# Analysis of leaf CO_2_ Assimilation, CO and CH_4_ Release Under Different Environmental Settings

**DOI:** 10.1101/2025.04.30.651537

**Authors:** Duarte Casanova, Dan Bruhn, Teis Mikkelsen

**Author notes:** These authors contributed equally to this work (Dan Bruhn, Teis Mikkelsen).

## Abstract

Many studies have found plant leaves to be emitters of CO and CH_4_. Consensus indicates that CH_4_ emissions are stimulated by heat and UV, while CO release is additionally stimulated by visible light. The mechanisms producing these emissions are yet to be discovered. To get closer to finding these mechanisms, this study examined whether photosynthesis might influence CO and CH_4_ leaf emissions. Five plant species of different photosynthesis pathways were analysed for their photosynthesis performance, as well as CO and CH_4_ emissions under different temperatures and visible light intensities. Findings reveal CO release rates to be positively correlated with light intensity and temperature but suggest a separate dark metabolism. CH_4_ rates were independent of light intensity and temperature. Much lower CH_4_ release from excised leaves compared to their connected counterparts, indicates that such is dependent on stomatal opening, supporting the hypothesis that CH_4_ is dissolved in transpired water. CO release rates are similar between attached and detached leaves, suggesting that CO is produced at epidermal level. Photosynthesis appears to be unrelated to the release of either of these gases.

**Key Message:** No link was found between CO and CH_4_ emission rates and CO_2_ Assimilation. Combining the results from this study with previous research, CO is concluded to be produced in the epidermis and CH_4_ to be dissolved in transpired water.

## 1 Introduction

Photosynthesis is the biological process through which most plants convert light energy into chemical energy. It involves the capture of carbon dioxide (CO_2_) and release of oxygen (O_2_), through epidermal pores called stomata. Photosynthetic Photon Flux Density (PPFD), refers to the number of photons within the light waveband usable for photosynthesis that reach a surface per unit of time.

In environments that may constrain photosynthesis, different plant species have adaptations that allow them to maintain high production rates. These are called photosynthetic pathways.

The reason plants have evolved to use different photosynthesis pathways is to deal with photorespiration. RubisCO, the same enzyme that initiates the Calvin Cycle, also commences a series of reactions, known as photorespiration, that result in the loss of 3 fixed carbons. This process is stimulated by higher temperatures and lower CO_2_:O_2_ concentration ratio (Greer, 2015).

The classic photosynthesis pathway, used by about 85-95% of the Earth’s plant species (Yamori et al., 2014; Boretti and Florentine, 2019), is known as C_3_. In arid conditions, C_3_ plants close their stomata to reduce transpiration losses. But this also reduces CO_2_ intake and O_2_ outflow, thus decreasing the CO_2_ : O_2_ ratio inside the leaf and increasing photorespiration. These environments have driven the evolutionary divergence into new photosynthesis pathways that can avoid high photorespiration rates.

In another pathway, known as C_4_, CO_2_ is initially fixed by the enzyme PEP carboxylase into a four-carbon compound, which is then transported to a bundle sheath cell. There, it is decarboxylated to release CO_2_ for use in the Calvin Cycle (Osborne and Beerling, 2006). This whole process involves a series of reactions that incur an extra energetic cost of 1-3 ATP per CO_2_ molecule transported (Nobel, 1991).

Some plants use the crassulacean acid metabolism (CAM) pathway to deal with dry environments. CAM plants open their stomata during the night, letting CO_2_ diffuse in and be fixed by PEP carboxylase, following a series of reactions to malic acid, which is stored inside vacuoles. During the day, CAM plants close their stomata to avoid losing water and CO_2_ is gradually released from the vacuoles into the Calvin Cycle. The CAM pathway also comes at a cost of 2-4 additional ATP for each CO_2_ molecule fixed (Nobel, 1991).

These physiological differences between plants can be seen at ecosystem level. C_3_ plants dominate in cold and moist environments, while C_4_ and CAM plants are mostly found in hot and dry ecosystems (Larcher, 2003), where their adaptations to reduce photorespiration are worth the added ATP costs.

Alongside photosynthesis, other unidentified leaf processes have been found to release carbon monoxide (CO) and methane (CH_4_).

Studies have shown that plant leaves release CO in the presence of light and oxygen. Bruhn et al. (2013) found a positive correlation between CO release and increases in temperature and UV intensity across multiple plant species. Irradiation appears to be more significant than temperature in driving CO emissions (Schade et al., 1999). Visible light accounts for up to 40% of the UV-induced CO emissions, with lower wavelengths inducing a higher release of CO (Schade et al., 1999). Another topic of interest is where this CO comes from. Schade et al. (1999) suggested that it may stem from cleavage of the cellulose chain. Wilks (1959) analysed chlorophyll extracts exposed to light, finding that the same component of the light spectrum absorbed in photosynthesis (480 to 680 nm) results in release of CO.

Recent observations of CH_4_ release by eukaryotes under aerobic conditions have lead researchers to try to identify the processes and conditions governing this release, and its contribution to the atmospheric methane budget. It is estimated that aerobic emissions provide 1.7-11.5% of the CH_4_ released into the troposphere (Kirschbaum et al., 2006; Parsons et al., 2006; Butenhoff and Khalil, 2007).

One major challenge in scaling up oxic CH_4_ release by plants is the uncertainty of results obtained from studies at the ecosystem level, as an unknown amount of the CH_4_ being produced comes from soil bacteria. Zeikus and Ward (1974) suggested that some observed CH_4_ release in attached leaves is actually soil-produced, being transported via the transpirational stream. This hypothesis is corroborated by Qaderi and Reid (2009), who found higher rates of CH_4_ release by attached pea leaves compared to detached ones.

Bruhn et al. (2012) documented the 3 most generally accepted factors to stimulate plant CH_4_ release to be increases in temperature, UV radiation and reactive oxygen species. The latter is a byproduct of other stresses, such as heat, UV and wounds. This led the authors to call reactive oxygen species a “ unifying mechanism for plant CH_4_ production” .

Nisbet et al. (2009) tried to map the pathway that conducts the CH_4_ release. They found that the same species of plant grown in different locations demonstrates notably different *δ*^13^C profiles in the emitted CH_4_. The authors therefore raise doubts about a plant-specific process for CH_4_ production, which they argue should result in a very similar isotopic composition. The authors conclude that the observed CH_4_ release is dissolved in transpired water.

## 2 Methodology

A series of experiments was conducted to try to uncover the yet-to-discover mechanisms underlying the leaf-release of CO and CH_4_.

Given that photosynthesis involves many leaf-scale processes and organelles, it may be directly or indirectly involved in the release of CO and/or CH_4_ from leaves. To investigate the correlation between photosynthesis and the release of these gases, different plant species’ photosynthesis rates were metered under various light levels and temperatures. CO and CH_4_ release rates were also calculated for the same conditions.

For measuring photosynthesis a LI-COR^®^ LI-6800 Portable Photosynthesis System was used. This device pumps air and measures its parameters before and after crossing a leaf sample inside a chamber. Reference air CO_2_ concentration ([CO_2_]), temperature (T) and relative humidity (RH) were controlled to be constant for the entirety of a set of measurements, making the loop an open system. In other words, for every circulation loop the air does, its parameters are reset before reaching the leaf, so the difference between the measured parameters before and after the leaf represents plant gas exchange.

Each leaf was tested under the temperatures of 15, 25 and 35^*°*^C. For each temperature, 13 different levels of PPFD (0, 100, 200, 300, 400, 500, 600, 700, 800, 900, 1000, 1500, 2000 *µ* mol · m^*−*2^ s^*−*1^) were applied to the leaf. The light used consisted of a spectral blend of 10% 453nm + 90% 660nm, especially targeting chlorophyll pigments.

Table 1 parameters were kept constant in all LI-6800 experiments:

**Table 1.** Constant parameters used in LI-6800 experiments.

| Constant | Value |
| --- | --- |
| % O<sub>2</sub> | 21% |
| [CO<sub>2</sub>] | 430 ppm |
| RH | 50% |
| Flow | 400 $\mu\text{mol} \cdot \text{s}^{-1}$ |

CO_2_ Assimilation (*A*) was used as a metric for photosynthesis. *A* is computed as *A* = *µ*(*c*_1_*− c*_2_)*/S* where *µ* represents the flow, *S* is the leaf area inside the chamber (2 cm^2^) and *c*_1_ and *c*_2_ are the CO_2_ concentrations before and after the sample, respectively.

After recording photosynthesis rates, the leaves were excised, washed with air depleted in CO and CH_4_, and placed inside a WALZ^®^ 3010-GWK1 Gas-Exchange Chamber connected to a Picarro^®^ G2401 Gas Concentration Analyzer. Unlike the LI-6800, this setup is a closed-loop system, meaning that the atmospheric parameters inside the setup continuously change. The advantage of a closed-loop is that it makes it possible to see small changes in trace gases such as CO and CH_4_.

The G2401 continuously logs the air concentrations of CO and CH_4_. Emission rates are obtained by calculating the linear regression slope of the gases’ concentrations over the 5-minute period following closure of the chamber.

Each leaf sample was exposed to the chamber temperatures of 15, 25 and 35^*°*^C. For each temperature, 6 different levels of PPFD (0, 100, 200, 600, 1000, 2000 *µ* mol *·* m^*−*2^ s^*−*1^) were applied to the chamber, using the same spectral blend as in the LI-6800. The light was emitted by a Heliospectra^®^ DYNA lamp and measured using a LI-COR^®^ LI-180 Spectrometer. During data collection, the setup was covered with reflective material to prevent external light from entering the chamber. For dark measurements, a black cloth was placed on top of the chamber to further block light.

In all measurements, a fan inside the chamber ensured continuous air mixing. Parameters such as RH were not controlled in this setup. CO and CH_4_ concentrations are corrected for H_2_O interference.

The gas release rates are obtained in g/h and are divided by the dry weight of the leaf sample, obtained by drying out the leaves in a 80^*°*^C oven for 48h.

Alternatively to the dry weight, *m*? is also divided by the leaf area, obtained by analysing leaf photographs with the software ImageJ.

Table 2 shows the plant species that were conducted tests on. Plants were chosen based on their photosynthesis pathway, origin and fitness of the leaves for the available equipment.

**Table 2.** Different plant species used, together with photosynthesis pathway, environmental temperature they were grown at (T) and number of replicates (N)

| Photosynthesis Pathway | Species | Origin | T (°C) | N |
| --- | --- | --- | --- | --- |
| C<sub>3</sub> | *Brugmansia sanguinea* | Andes | 12 | 5 |
| C<sub>3</sub> | *Coffea arabica* | Arabia | 16 | 2 |
|  |  | Ethiopia | 12 | 2 |
| C<sub>4</sub> | *Saccharum officinarum* | Saint Croix | 16 | 5 |
| CAM | *Kalanchoe fedtschenkoi* | Madagascar | 18.5 | 5 |
| CAM | *Vanilla planifolia* | Mexico | 21 | 5 |

Table 2 shows the plant species used.

Each replicate corresponds to a different leaf. For all plants except *Coffea arabica*, all individuals used were clones. To maximize behavioral variation, the replicates are either leaves from different heights of the same plant or from genetically-identical clones.

Given the consistency of photosynthesis rates across replicates, only 3 replicates were used per plant in the LI-6800 measurements.

All plants used were grown in the University of Copenhagen Botanical Garden in Gothersgade 128, under different environmental conditions. The time it took from leaf excision to completion of experiments was no longer than 4 hours.

## 3 Results and Discussion

### 3.1 C_3_ and C_4_ Photosynthesis

Photosynthetic rates can serve as indicators of plant adaptation to different environments. A particularly interesting case is comparing how C_3_ and C_4_ plants’ assimilation curves respond to increasing temperatures.

C_4_ plants possess adaptations to minimize photorespiration, which can become prominent in the C_3_ pathway under high temperatures. As a result, in higher temperatures C_3_ plants may demonstrate lower assimilation rates as more RubisCO will bind to O_2_ instead of CO_2_ (Ghashghaie and Cornic, 1994).

Indeed, a slight depression of photosynthesis was observed when temperature was increased from 25 to 35^*°*^C in the C_3_ *Brugmansia sanguinea*. Such did not occur in the C_4_ plant studied, as seen in Fig. 1.

**Fig. 1.**
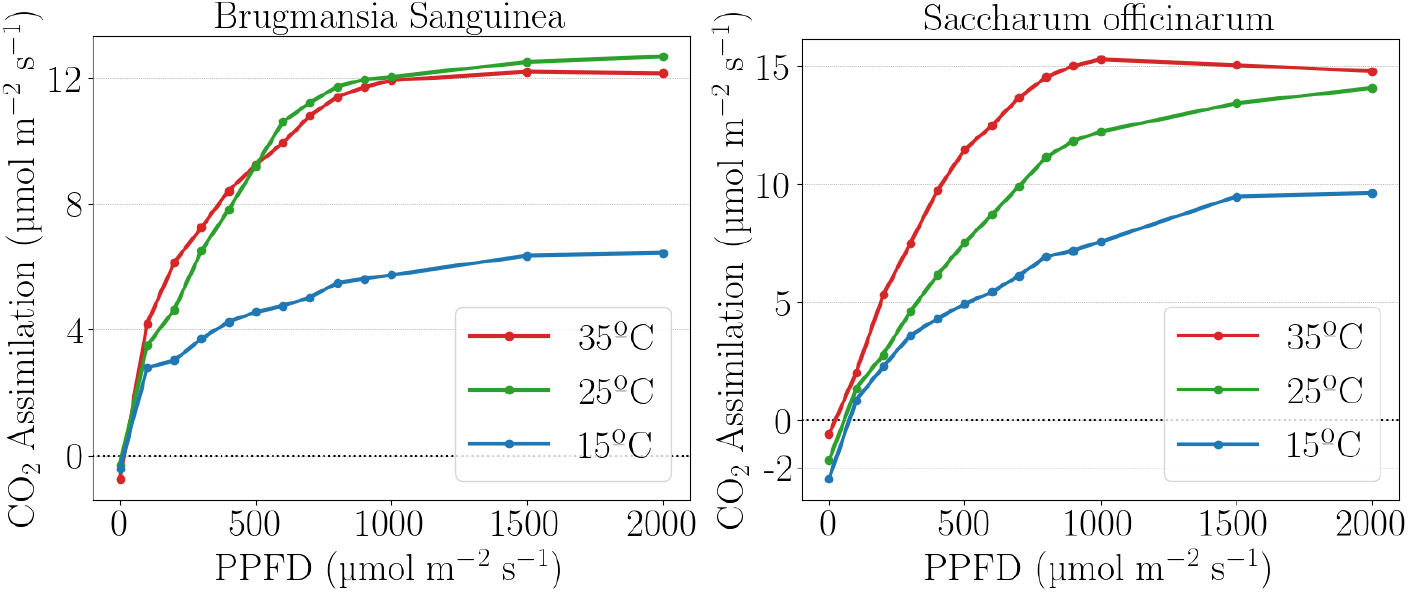
CO_2_ uptake for different levels of PPFD for a) *Brugmansia sanguinea*, a C_3_ plant, and b) *Saccharum officinarum*, a C_4_ plant

These results aline with Yamori et al. (2014)’s claim that the optimal temperature for photosynthesis is higher in C_4_ than C_3_ plants.

Notably in all C_3_ and C_4_ plants tested, assimilation decreased past PPFD=1000 *µ* mol · m^*−*2^ s^*−*1^ when *T* = 35^*°*^*C*. In the case of *Coffea arabica*, a C_3_ plant, assimilation rates were highest at 35^*°*^C and began to decrease from PPFD levels as low as 500 *µ* mol · m^*−*2^ s^*−*1^.

The negative assimilation values seen in darkness are exhibitions of respiration, as the absence of light halts photosynthesis. In the majority of replicates, respiration in darkness was higher the higher the temperature.

### 3.2 CAM Photosynthesis

Overall, lower rates of photosynthesis were observed in CAM plants. In a few replicates, CAM leaves produced a photosynthesis curve like that of a C_3_/C_4_ plant, exhibiting negative assimilation in darkness and logarithmically increasing their assimilation with light. This may corroborate the hypothesis that CAM plants can switch to a C_3_/C_4_-like style of photosynthetic metabolism when circumstances are favorable (Osmond et al., 1973).

Different replicates of both CAM plants exhibited very different photosynthesis profiles. This indicates that the temporal duality of the photosynthetic process in CAM plants is governed by factors not manipulated in this study. Figure 2 shows the photosynthesis rates of different replicates under the same conditions. Turquoise represents the 1^*st*^, purple the 2^*nd*^, and pink the 3^*rd*^ replicate.

**Fig. 2.**
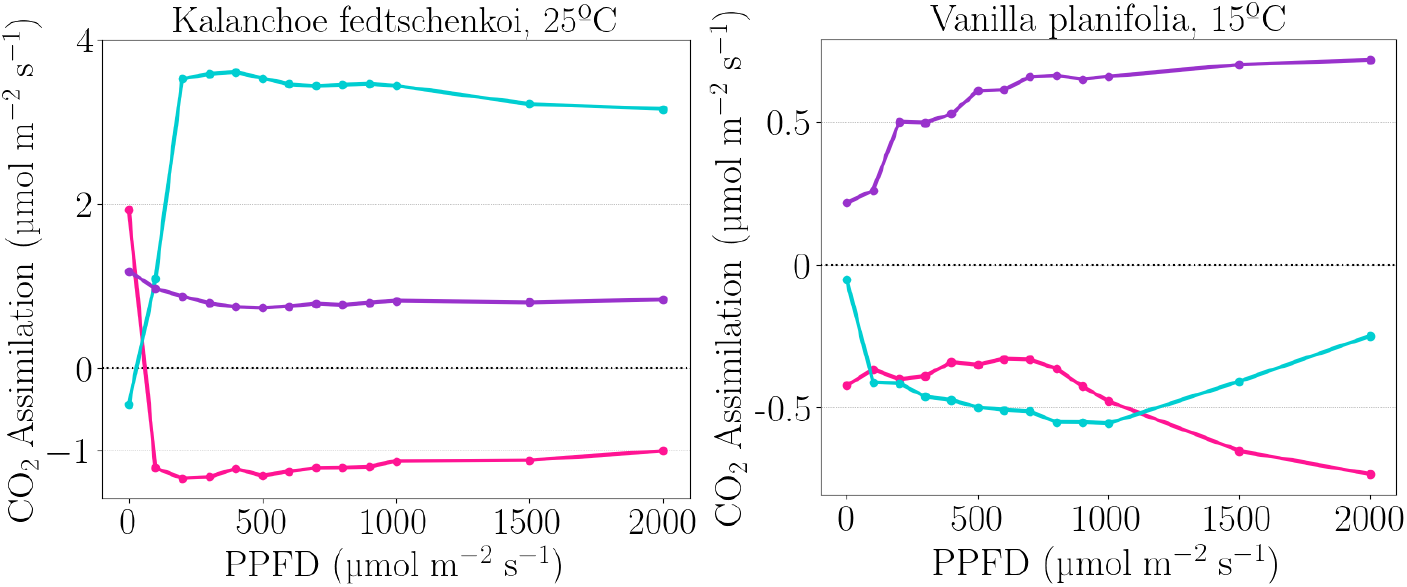
CO_2_ uptake for different levels of PPFD for 3 different replicates of a) *Kalanchoe fedtschenkoi* at 25^*°*^C and b) *Vanilla planifolia* at 15^*°*^C

Given the inconsistent photosynthesis curves obtained from CAM plants, it was opted not to treat the different replicates together.

#### 3.2.1 Stomatal Opening Controls

The different photosynthesis curves for each replicate raise the question of what regulates stomatal opening in CAM plants.

To examine if the temporal duality of the CAM process is the reason behind the different patterns across replicates, tables 3 and 4 categorize each measurement according to the time of the day.

**Table 3.** *Kalanchoe fedtschenkoi* replicates.

| Replicate | Day | Time |
| --- | --- | --- |
| 1 | 10/5 | 8:00 |
| 2 | 8/6 | 10:15 |
| 3 | 8/6 | 12:15 |

**Table 4.** *Vanilla planifolia* replicates.

| Replicate | Day | Time |
| --- | --- | --- |
| 1 | 1/6 | 9:20 |
| 2 | 5/6 | 10:10 |
| 3 | 7/6 | 11:00 |

The differences in temporal order do not explain the different behaviours between replicates, suggesting that the photosynthetic behavior of a CAM plant isn’t based on time of the day.

Remarkably, CO_2_ uptake increased with temperature for the majority of replicates. The average CO_2_ assimilation across different light levels shows a positive correlation with temperature, as seen in Fig. 3.

**Fig. 3.**
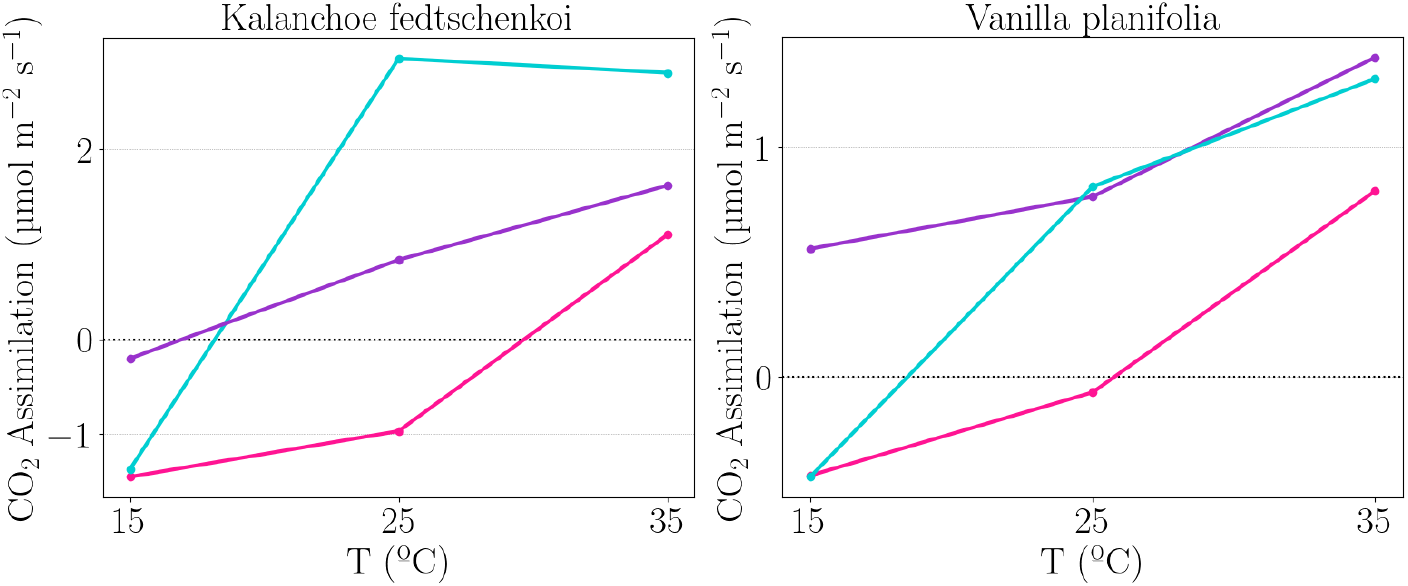
Average CO_2_ uptake across all levels of PPFD for 3 different replicates of a) *Kalanchoe fedtschenkoi* and b) *Vanilla planifolia*, according to temperature

Given that the relative humidity was kept constant at 50% for all measurements, the increases in assimilation with temperature indicate that stomatal closure to prevent water loss is not triggered by temperature changes. Instead, the stomatal closure and opening mechanism must be regulated by another factor. Dayanandan and Kaufman (1975) observed the stomata of *Kalanchoe fedtschenkoi* and other CAM plants and reported that potassium ions migrate in and out of guard cells, creating turgor pressure that opens and closes them.

However, recent research has shown that potassium ion migration isn’t enough to create turgor pressure (Lee, 2010). During the night CAM plants produce malic acid which builds turgor pressure inside guard cells, leading them to open (Kim and Lee, 2007). Through daytime, malic acid is broken down to obtain CO_2_ for the Calvin cycle, dropping the turgor pressure and closing the stomata.

This way the stomatal aperture cycle follows the circadian rhythm and is controlled not only by the intensity of the light, but also by potassium and malic acid concentrations inside guard cell vacuoles. This explains why different replicates behaved differently in CAM plants, as intracellular factors were not the controlled in this experiment.

### 3.3 CO

Results demonstrate plants release CO at all temperatures and light intensities, as seen in Figures 4 and 5.

**Fig. 4.**
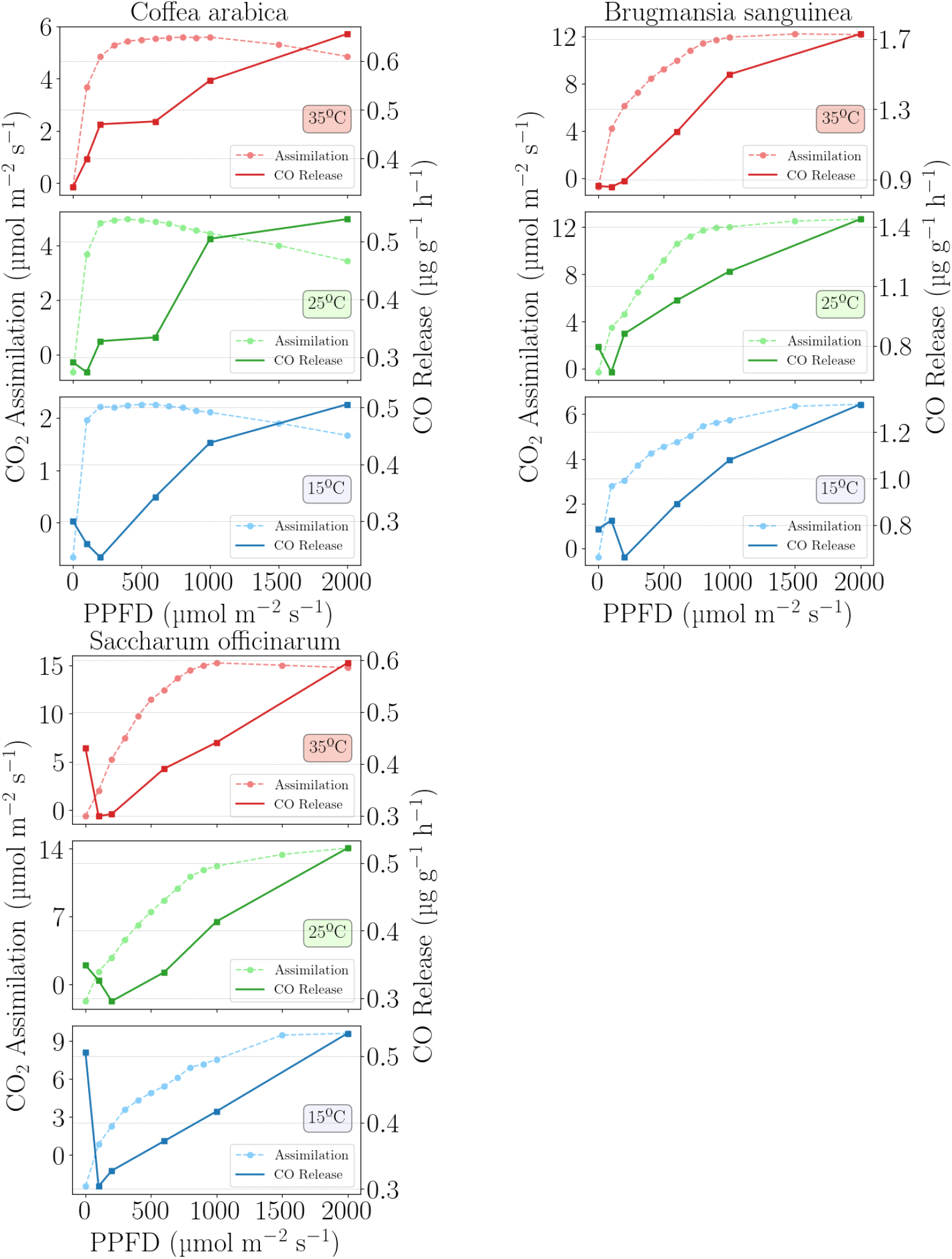
CO_2_ assimilation (dashed lines) and CO release (solid lines) for different levels of PPFD and temperatures for a) *Coffea arabica*, b) *Brugmansia sanguinea* and c) *Saccharum officinarum*. The magnitude in the y-axis varies between graphs to improve visibility

**Fig. 5.**
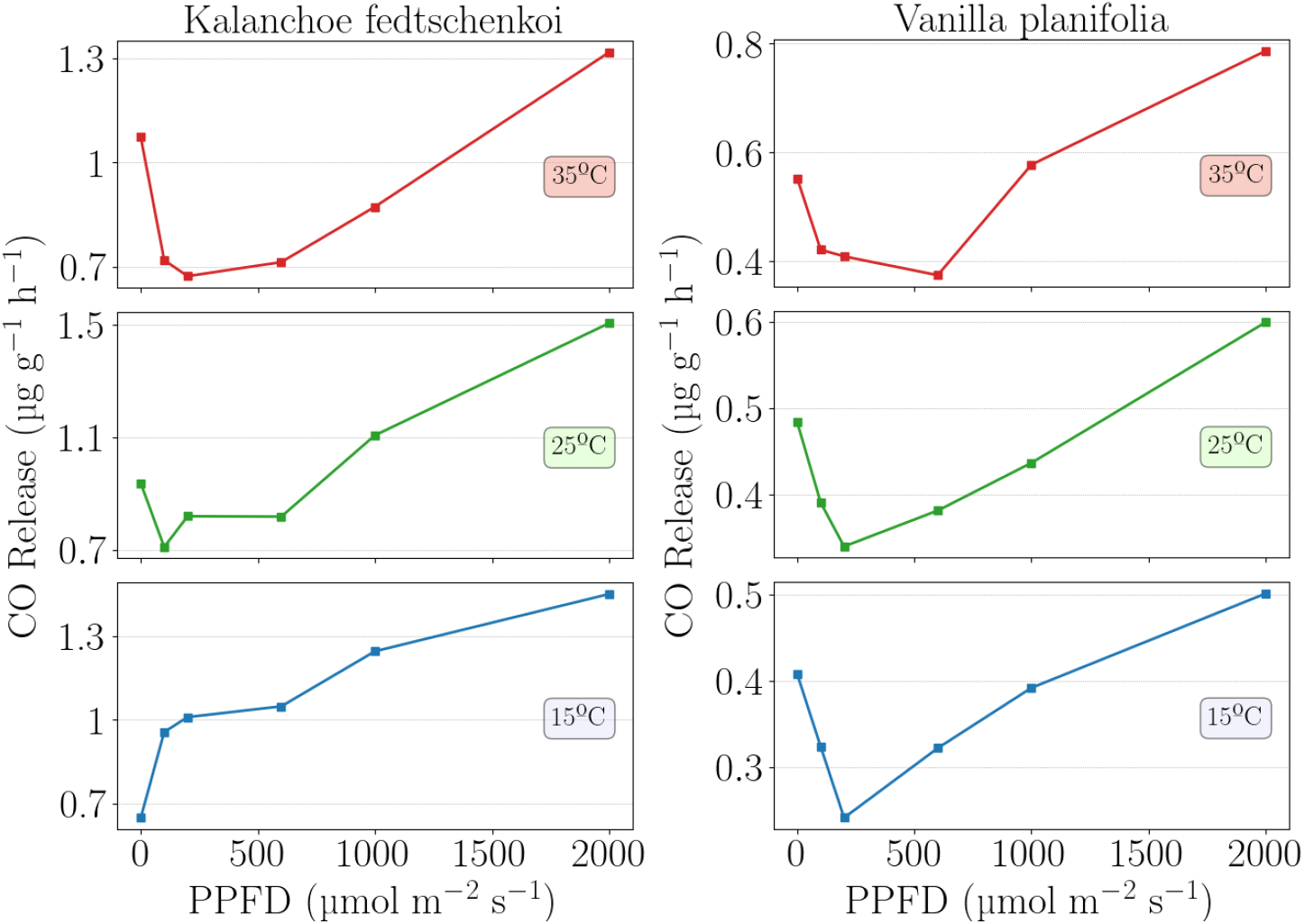
CO release for different levels of PPFD and temperatures for a) *Kalanchoe fedtschenkoi* and b) *Vanilla planifolia*. The magnitude in the y-axis varies between graphs to improve visibility

In C_3_ and C_4_ species, the CO rate generally increases with light intensity, often in a linear fashion, consistent with observations by Seiler et al. (1978) and Bauer et al. (1979) in tree leaves.

For some species, darkness induced more CO release than low light levels. This is particularly remarkable in *Vanilla planifolia* which produced a U-shaped emission curve regardless of temperature. Moreover, C_3_/C_4_ plants’ darkness measurements often produce rates outside the linear trend of light measurements, indicating that there may be separate dark metabolic pathways producing CO. These were most noticeable on *Saccharum officinarum*, the only C_4_ plant analysed. The observed CO release in darkness disputes Wilks (1959)’s suggestion that such requires both oxygen and light.

In general, increases in temperature led to higher CO emissions from leaves, as well as wider emission ranges depending on irradiation.

The relation between CO release and photosynthesis remains undisclosed. Like photosynthesis, CO release is stimulated by visible light. But unlike photosynthesis it is not reliant on it, and CO rates didn’t show signs of capping at high light intensities. Moreover, the sensored CO emissions may not correspond exactly to the CO released from the leaves. Logan et al. (1981) claim that an unknown, high share of the CO observed in experiments is actually produced by the oxidation of hydrocarbons released by plants, such as isoprene (Zimmerman et al., 1978). This might be particularly noteworthy in closed-loop system measurements, such as this one.

Emission rates varied considerably from replicate to replicate, producing high errors for many of the measurements.

### 3.4 CH_4_

Results demonstrate plants release CH_4_ at all temperatures and light intensities, as seen in Figures 6 and 7.

**Fig. 6.**
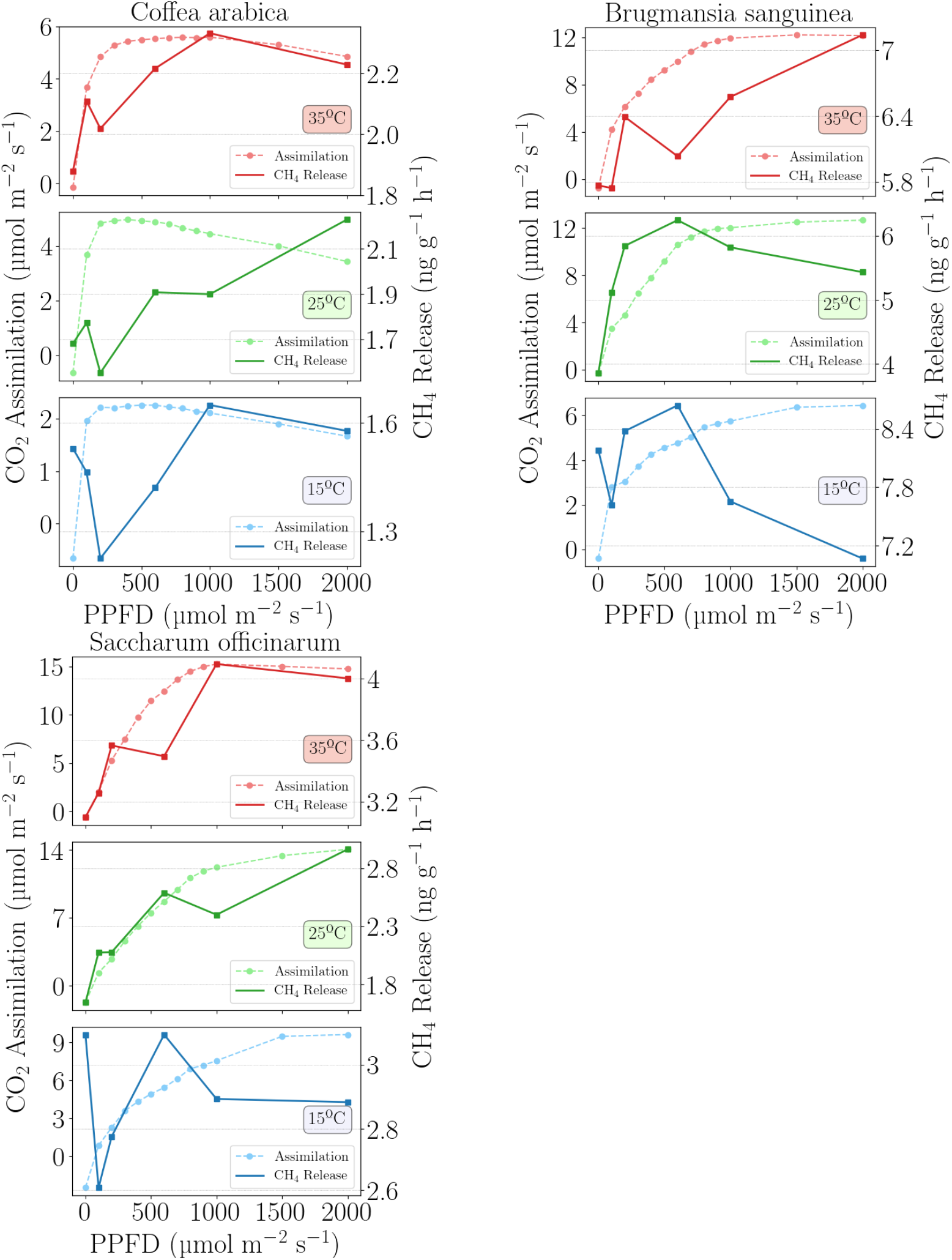
CO_2_ assimilation (dashed lines) and CH_4_ release (solid lines) for different levels of PPFD and temperatures for a) *Coffea arabica*, b) *Brugmansia sanguinea* and c) *Saccharum officinarum*. The magnitude in the y-axis varies between graphs to improve visibility

**Fig. 7.**
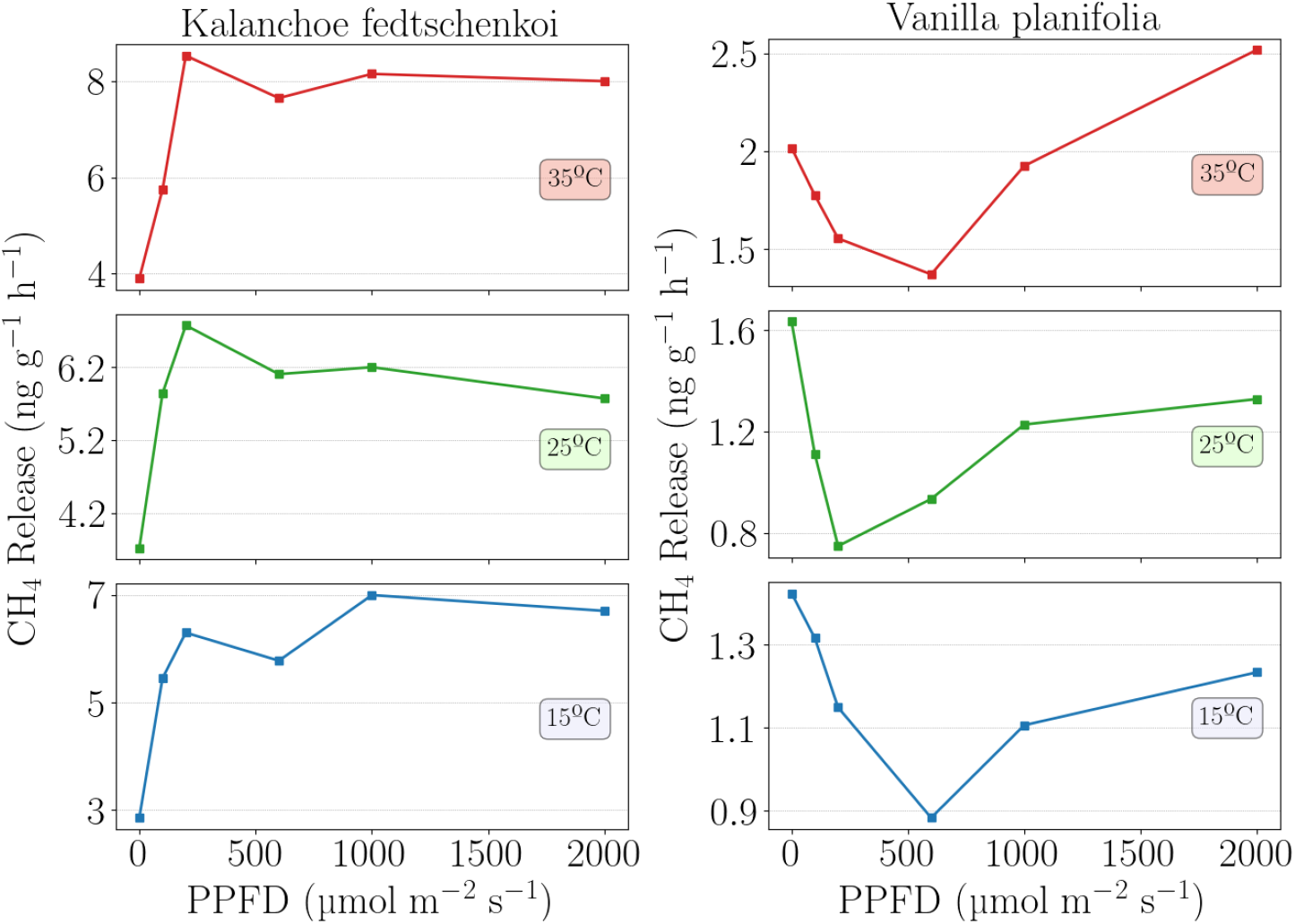
CH_4_ release for different levels of PPFD and temperatures for a) *Kalanchoe fedtschenkoi* and b) *Vanilla planifolia*. The magnitude in the y-axis varies between graphs to improve visibility

CH_4_ emission patterns are more chaotic than those of CO. Unlike CO, an increase in temperature doesn’t translate into an increase in CH_4_ emissions for some of the plants tested. The range of emissions seems independent from temperature as well. Singularly, CH_4_ emissions didn’t show an exponential correlation with temperature, as observed in previous studies (Keppler et al., 2006; Vigano et al., 2008).

Among the CAM plants tested, a discernible pattern emerged across the different temperatures. For *Kalanchoe fedtschenkoi* there is a logarithmic increase in emissions followed by a slight decrease beyond PPFD=1000, in a fashion similar to that of photosynthesis curves of C_3_/C_4_ plants. For *Vanilla planifolia* the U-shape is once again observed, as with CO, suggesting that both gases may originate in the same biochemical process.

The magnitude of CH_4_ emissions observed in this study is way lower than that of previous studies on attached leaves. Keppler et al. (2006) recorded CH_4_ emissions for attached leaves to be on the range 198-598 *ng* · g^*−*1^ h^*−*1^. However, for excised leaves the authors found a much lower range of emissions, 1.6-15.8. Nisbet et al. (2009) found a similar range of 1-20 for detached leaves.

In this investigation, excised leaves were used to measure CH_4_ release and an emission range of 0.9-8.5 ng · g^*−*1^ h^*−*1^ was obtained, matching the measurements of Keppler et al. (2006) and Nisbet et al. (2009).

## 4 Possible Origins of CO and CH_4_

Combining the findings from this study with existing research, it is possible to draw theories for the potential origins of CO and CH_4_.

### 4.0.1 CH_4_

The most protruding feature about plant CH_4_ emission is the gap between connected and excised leaves. The reason for this may be stomatal activity.

When leaves are excised, stomatal aperture falls rapidly (McAdam and Brodribb, 2012). As a result, the CO and CH_4_ measurements presented in this report represent the gas releases by leaves with partially closed stomata.

The remarkable difference between CH_4_ release from connected leaves and their excised counterparts may stem from the difference in stomatal aperture. Given that connected leaves have CH_4_ release rates of much higher magnitude, it is speculated that CH_4_ is mostly being released through the stomata.

The hypothesis that CH_4_ is being released through the stomata corroborates Zeikus and Ward (1974) and Nisbet et al. (2009)’s suggestion that CH_4_ is produced in soil and transported through the xylem.

The findings of Vigano et al. (2008) conclude that UV incidence on excised leaves increases CH_4_ formation. This may be explained by increased transpiration caused by UV light (Phyo and Chung, 2013), or indicate towards an independent leaf CH_4_ production mechanism that is stimulated by UV (or its by-products, such as free radicals).

### 4.0.2 CO

Compared to CH_4_, the difference in CO emissions from excised and connected leaves is less pronounced: Bruhn et al. (2013) calculated CO rates of 2466 nmol m^*−*2^h^*−*1^ for a grass field and 1740 nmol m^*−*2^h^*−*1^ for freshly cut leaves (using sunlight, excluding most UV).

This indicates that CO release is mostly independent from stomatal aperture. As such, CO may instead originate from processes occurring at the leaf’s surface.

The leaf epidermis is covered by a film known as cuticle. The latter is enveloped by waxes, composed of a mixture of alkanes, alcohols, ketones and esters (Kunst and Samuels, 2009). These compounds may be subject to reactions with surrounding atmospheric gases and produce CO. The reactions causing release of CO are expected to be stimulated by high temperatures and visible light, as was observed in this study.

The reason why dark measurements sometimes demonstrated higher CO release than light measurements remains unresolved. It points towards an unknown, CO-producing, dark metabolism that is diminished by light. Additional research is needed to clarify dark CO release, as the positive rates found in this study contradict Bruhn et al. (2013)’s findings of null CO release in darkness, despite the authors’ similar CO rates for light measurements.

In summary, it is speculated that a high share of the CH_4_ measured in attached leaves is produced in the soil, and may be complemented with independent leaf CH_4_ production. The observed CO is likely being produced by reactions happening at the leaf’s surface.

## 5 Conclusions

It was found that in C_3_ and C_4_ plants, photosynthesis rates began to decline at high T (35^*°*^C) past PPFD=1000 *µ* mol · m^*−*2^ s^*−*1^ and that CAM photosynthesis benefits from increases in T. CO emissions increased linearly with PPFD, though darkness often results in higher emission rates than low PPFD. T also had a direct correlation with CO release. CH_4_ emissions on the other hand appear to be unrelated to both PPFD and T. CO emissions are suggested to be produced at epidermal level, given their low dependence on stomatal opening, while CH_4_ is believed to be dissolved in transpired water, but may also be produced by the plant. No link was found between photosynthesis curves and release of CO and CH_4_.

## Aknowledgements

We acknowledge the help of the Copenhagen Botanical Garden in providing leaf samples and environmental data.

